# Comparison of adopted and non-adopted individuals reveals gene-environment interplay for education in the UK Biobank

**DOI:** 10.1101/707695

**Authors:** Rosa Cheesman, Avina Hunjan, Jonathan R. I. Coleman, Yasmin Ahmadzadeh, Robert Plomin, Tom A. McAdams, Thalia C. Eley, Gerome Breen

## Abstract

Individual-level polygenic scores can now explain ∼10% of the variation in number of years of completed education. However, associations between polygenic scores and education capture not only genetic propensity but information about the environment that individuals are exposed to. This is because individuals passively inherit effects of parental genotypes, since their parents typically also provide the rearing environment. In other words, the strong correlation between offspring and parent genotypes results in an association between the offspring genotypes and the rearing environment. This is termed passive gene-environment correlation. We present an approach to test for the extent of passive gene-environment correlation for education without requiring intergenerational data. Specifically, we use information from 6311 individuals in the UK Biobank who were adopted in childhood to compare genetic influence on education between adoptees and non-adopted individuals. Adoptees’ rearing environments are less correlated with their genotypes, because they do not share genes with their adoptive parents. We find that polygenic scores are twice as predictive of years of education in non-adopted individuals compared to adoptees (R^2^= 0.074 vs 0.037, difference test p= 8.23 × 10^−24^). We provide another kind of evidence for the influence of parental behaviour on offspring education: individuals in the lowest decile of education polygenic score attain significantly more education if they are adopted, possibly due to educationally supportive adoptive environments. Overall, these results suggest that genetic influences on education are mediated via the home environment. As such, polygenic prediction of educational attainment represents gene-environment correlations just as much as it represents direct genetic effects.

## Introduction

An important process by which genes and environments work together to influence behaviour is gene-environment correlation. Gene-environment correlation refers to the association between the genotype an individual inherits from their parents and the environment in which they are raised (Plomin et al. 1977). In behaviour genetics, three forms of gene-environment correlation are distinguished: passive, active and evocative. An example of passive gene-environment correlation is that more educated parents are likely to provide both beneficial genes and educationally supportive family environments, such as books in the home, for their children. Therefore, shared genes confound associations between putative environmental variables and child attainment. This phenomenon is of interest across disciplines and is also known as ‘dynastic effects’, ‘genetic nurture’, and ‘social genetic effects’. Active and evocative gene-environment correlations reflect how genotypes lead to phenotypes: individuals select and evoke environments based on their genetically influenced traits.

It is essential to investigate gene-environment interplay in educational attainment, for several reasons. First, educational attainment is an important trait for individuals and society. Second, it is clear that gene-environment correlation matters for educational attainment. Adoption, twin, and instrumental variable research suggests that shared genes largely explain associations between parent and child educational attainment (Holmlund et al. 2011). Third, genome-wide polygenic scores, which index the genetic liability that each individual carries for a specific trait, are notably powerful for education attainment and now predict ∼10% of the variation in years of education (Lee et al. 2018), with potential social implications (Plomin and von Stumm 2018). However, this prediction may be influenced by both direct genetic effects on an individual’s own education and passive gene-environment correlation. Since genetic variants present in offspring are also present in their parents, they can also have indirect effects on offspring education via the parents’ genetically influenced behaviour. It has only recently been shown that, through this indirect pathway of passive gene-environment correlation, offspring polygenic scores partially index the family environment (Kong et al. 2018; Bates et al. 2018).

Adoption studies have been key in attempts to disentangle the causal processes affecting educational attainment. Adoption designs remove the overlap between genetic and environmental influences (passive gene-environment correlation). This is achieved by measuring the resemblance of adopted children with their birth parents, who are genetically related but do not share an environment, and with their adoptive parents, who are genetically unrelated but share an environment. The former gives an estimate of direct genetic influence, independent of passively correlated environmental effects, whereas the latter gives an estimate of shared environmental influence, free of correlated genetic effects. It is also possible to estimate passive gene-environment correlation as the extent to which genes contribute more to the covariation between measures of the family environment and offspring traits in non-adoptive than adoptive families (Plomin et al. 1985). Notably, other forms of gene-environment correlation are still present in adoptees, since heritable proclivities lead them to select and evoke experiences.

More recently, researchers have applied genomic tools to family data to estimate direct and indirect effects on educational attainment (Bates et al. 2018; Kong et al. 2018; Domingue et al. 2015; Wertz et al. 2018; Selzam et al. 2019; Young et al. 2018). These designs can be thought of as conceptually related to adoption designs, since they account for shared genes between parents and offspring. Many of these studies have used polygenic scores. Passive gene-environment correlation can be estimated by creating polygenic scores for genetic variants that were not passed on by parents, and thus can only have indirect effects on offspring traits, through genetically-influenced parental behaviour (Bates et al. 2018; Kong et al. 2018). For example, using an educational attainment polygenic score based on non-transmitted genetic variants to control for indirect effects, the variance explained by the transmitted score shrank from 5 to 2% (Kong et al. 2018). The non-transmitted score also independently predicted attainment. The family environment is an important contributor to polygenic score prediction because it is adding to estimates of genetic influence, and because parents still influence their offspring after controlling for shared (transmitted) genes.

Our main aim was to use the ‘natural experiment’ created by adoptive placement to measure the importance of passive gene-environment correlation for educational attainment. When children are adopted by non-relatives, the indirect genetic path between the rearing environment and their traits is not present because adoptive parents are not genetically related to adopted children. This leads to three hypotheses. First, the phenotypic variance is expected to be lower in adopted individuals compared to non-adopted individuals. This is because adoptees do not have the additional source of variance that is due to passive gene-environment correlation (Loehlin & De Fries 1987; Plomin 1994). It could also be because adoptive families may be similar in their education level and socio-economic status or may be selected for better perceived parenting ability than is the average in non-adoptive families (Rutter 2006; Natsuaki et al. 2019). Second, if passive gene-environment correlation inflates standard estimates of heritability, then heritability should be lower in adopted individuals than in non-adopted individuals, because adopted individuals are reared in environments that are less correlated with their genotypes. Third, for the same reason, the variance explained by polygenic scores will be lower in adopted individuals, and may thus be closer to the true direct genetic effect of an individual’s own DNA. To shed further light on the moderation of polygenic prediction by adoption status, we also tested for gene-environment interaction using a model with polygenic scores for education as the predictor, adoption as the exposure and education as the outcome. As a negative control, we tested the polygenic prediction comparison between adoptees and non-adopted individuals for height, which has not shown evidence of passive gene-environment correlation in previous studies (Kong et al. 2018; Selzam et al. 2019).

The experience of adoption is unusual and may carry stressors. This could lead adoptees to differ systematically from non-adopted individuals. We therefore also assessed the comparability of the adopted and non-adopted groups. Specifically, we estimated the heritability of being an adoptee, and tested for differences in genetic effects on education in adopted versus non-adopted individuals. Lastly, we assessed the robustness of our results across different year-of-birth cohorts.

## Methods

### Sample, genotype quality control and phenotype definition

The UK Biobank is an epidemiological resource including British individuals aged 40 to 70 at recruitment (Allen et al. 2014). UK Biobank participants were asked “Were you adopted as a child?”. 8,040 individuals said yes, and 541,889 individuals said no. No additional information was collected on the age of adoption, or whether individuals were adopted by biological relatives. Genome-wide genetic data came from the full release of the UK Biobank data, and were collected and processed according to the quality control pipeline (Bycroft et al. 2018). We restricted analyses to individuals with full phenotypic data for education, who also passed genotype quality control criteria. This left 6,311 adopted and 375,343 non-adopted individuals for analysis.

Genotype quality control criteria were: common genetic variants of minor allele frequency > 0.01 that were directly genotyped or imputed with high confidence (INFO metric > 0.4); and individuals with genotype call rate > 98% who had concordant phenotypic and genetic gender information and who were unrelated to others in the dataset (less than third degree relatives). We performed removal of relatives using a “greedy” algorithm to minimise the exclusion of adoptees. To reduce confounding from population stratification, all analyses were limited to individuals of European ancestries, as defined by 4-means clustering on the first two genetic principal components provided by the UK Biobank. We also controlled for 10 ancestry principal components of the European sample in all genomic analyses.

Years of education, a proxy for educational attainment, was defined according to ISCED categories, as in previous genomic studies of the phenotype in UK Biobank and other samples (Lee et al. 2018). The response categories were: none of the above (no qualifications) = 7 years of education; Certificate of Secondary Education (CSEs) or equivalent = 10 years; O levels/GCSEs or equivalent = 10 years; A levels/AS levels or equivalent = 13 years; other professional qualification = 15 years; National Vocational Qualification (NVQ) or Higher National Diploma (HNC) or equivalent = 19 years; college or university degree = 20 years of education.

### Analyses

#### Phenotypic comparisons

First, we formally tested the hypothesis that non-adopted individuals show greater phenotypic variance than adopted individuals due to the presence of an additional source of variance (passive gene-environment correlation). A non-parametric test was used given the non-normal distribution of the education years variable (Brown and Forsythe 1974). This test is based on absolute deviations from the median, rather than the group mean. We also tested for differences in education years, age, and sex between the two groups, using a Wald test, z-test, and Wilcoxon test, respectively.

#### SNP heritability estimation

Second, to test the hypothesis that heritability is lower in adoptees, whose rearing environments are less correlated with their genotypes, we estimated the variance explained by common genetic variants for years of education in adoptees using Genomic-RElatedness-based restricted Maximum-Likelihood (GREML) (Yang et al. 2011), and compared this to the heritability estimate for non-adopted individuals. The method estimates heritability as the extent to which genetic similarity among unrelated individuals can predict their trait similarity. In GREML, a matrix of genomic similarity for each pair of unrelated individuals across genotyped variants is compared to a matrix of their pairwise phenotypic similarity using a random-effects mixed linear model, such that the variance of a trait can be decomposed into genetic and residual components, using maximum likelihood. We used two genetic relatedness matrices: one for adopted individuals, and a second for a subset of 6,500 non-adopted individuals. This was to enable comparison of two similarly sized samples, and to reduce the computational burden that results from scaling GREML to a sample as large as the UK Biobank. For both genomic matrices we used a relatedness cutoff of 0.025. Sub-samples were made using the “sample_n” function in the dplyr package in R (version 3.5). We compared these results to heritability estimates derived from a second method, LD score regression (LDSC) (B. K. Bulik-Sullivan et al. 2015). Unlike GREML, LDSC does not require individual-level data, allowing it to be computationally feasible to estimate the heritability of education in the full sample of non-adopted individuals. LDSC also enabled us to estimate genetic correlations (see below).

#### Polygenic scoring

Third, we tested whether the power of polygenic scores is greater for individuals who were reared with their biological relatives than for adoptees. The sample of non-adopted individuals was subdivided into three independent groups for polygenic score analyses. Our first sample consisted of 318,843 non-adopted individuals for genome-wide association analysis (GWA). The purpose of this was to estimate the effect sizes of associations between genome-wide genetic variants and years of education, to use for the creation of individual-level polygenic scores. We derived our base summary statistics file for years of education by meta-analysing summary statistics from our own GWA analysis in this subsample with independent summary statistics obtained from the Social Science Genomics Consortium (excluding UK Biobank and 23&Me) (Lee et al. 2018). The sample size for these external summary statistics was 324,160, leading to a total sample size of individuals in our GWA meta-analysis of 643,003.

Our second independent sample included 50,000 individuals to use for training our polygenic scores for years of education, i.e. identifying the optimal p-value threshold for inclusion of SNPs. The standard set of P-values in PRSice 2 were tested: 0.001, 0.05, 0.1, 0.2, 0.3, 0.4, 0.5, 1 (Choi Shing Wan n.d.).

Our third independent sample included 6,500 individuals, to match the sample size of adopted individuals, in which to run polygenic prediction models. In these prediction models we regressed the years of education phenotype in the UK Biobank on polygenic scores for years of education in adoptees, and then repeated the analysis in the 6,500 non-adopted individuals. In this third set of analyses we used a set p-value threshold obtained from the training step. This exact sample was the same as the one used to estimate SNP heritability of years of education. Notably, this polygenic score analysis is better-powered than the SNP heritability analysis, since it capitalises on the power of the large discovery sample (N=643,003).

As a negative control, we estimated the variance explained in height by a polygenic score for height (Wood et al. 2014) in adoptees versus non-adopted individuals. As with the education analysis, we trained the polygenic score in the sample of 50,000 individuals, and then tested the prediction at the best p-value threshold in our two independent and similarly sized samples of adopted and non-adopted individuals.

### Supplementary analyses

#### Heritability of adoption status

Substantial heritability of our environmental moderator might affect the interpretation of our main results. To explore this, we also estimated the heritability of adoption status using LD score regression in the full sample (N=381,654 individuals). The genetic ‘influence’ on adoption largely arises in the biological parent generation because heritable traits influence the likelihood of adoption of their child.

#### Polygenic score by adoption interaction analyses

Differences in genetic influences on the same trait across contexts – in this case adoption – can also be conceptualised as gene-environment interaction, whereby the impact of genes on educational attainment may be contingent on adoption status. We aimed to further explore our main results by testing a formal polygenic score by adoption interaction regression model. The model included main effects for polygenic score for years of education, adoption, and covariates, plus the interaction term as well as interaction terms for polygenic score and adoption with each covariate (Keller 2014). We tested a linear model for additive interaction and a logistic model for multiplicative interaction. To visualise any interaction, we plotted the regression slopes for polygenic prediction of educational attainment for adopted and non-adopted individuals (with both variables standardised to have a mean of 0 and a standard deviation of 1). Additionally, we stratified polygenic scores for adopted and non-adopted individuals overall (N=12,811) into deciles and tested for mean differences in years of education between adopted and non-adopted groups in each decile.

#### Qualitative differences in the genetic influence on education by adoption status

We assessed whether education is driven by the same set of genetic influences in adopted and non-adopted individuals. First, we estimated the genetic correlation between education in our samples. For this, we ran genome-wide association analyses of years of education in the full sample of non-adopted individuals (N= 375,343) and in the sample of adoptees, then estimated the genetic correlation between them using LD score regression. Second, we tested whether education is genetically linked to different traits between adoptees and non-adopted individuals. To this end, we estimated genetic correlations between education years and 247 traits available on LD Hub, for both adopted and non-adopted individuals. We compared the magnitudes of genetic correlations between education years and other variables between the adopted and non-adopted samples with z-tests.

#### Birth year-related differences in genetic influence

During the period when UK Biobank participants were growing up (1930s-70s), access to education increased, and there was great change in the norms and regulations surrounding reproduction, contraception and adoption. Previous studies have found that genetic influence on years of education increased in this period in the UK, since environmental differences between people had less influence on whether they stayed in education (Lee et al. 2018). We investigated temporal change in patterns of genetic influence on education in adopted versus non-adopted individuals by stratifying polygenic prediction analyses according to year of birth. Specifically, all individuals were split into 7 mutually exclusive birth-year groups, each with a range of 5 years, and polygenic score analyses were conducted separately for each of the year groups.

All analyses (SNP heritability, polygenic scoring, GWA) controlled for the following covariates: sex, age, 10 ancestry principal components, and factors capturing genotyping batch and centre. The majority of the analyses were completed in R version 3.5. GREML was performed in the GCTA software (Yang et al. 2011). Genome-wide association meta-analysis was performed in METAL (Willer et al. 2010). Polygenic score analyses were performed in PRSice 2 (Choi Shing Wan n.d.). To compare polygenic score results between adopted and non-adopted individuals, we obtained bootstrapped standard errors for the R^2^ statistics using the boot package in R, with 1000 replications. Genome-wide genetic correlations were estimated using LDSC (B. Bulik-Sullivan et al. 2015) and LD Hub (Zheng et al. 2017). The UK Biobank is a controlled-access public dataset available to all bona fide researchers.

## Results

### Sample analysed

The total sample of individuals with education phenotype data and quality-controlled genotype data was 381,654. As described in the Methods, individuals were split into 4 mutually-exclusive groups: a) adopted as children (N=6,311), b) 318,843 non-adopted individuals for genome-wide association analysis, c) 50,000 non-adopted individuals for training of polygenic scores, and d) 6,500 non-adopted individuals for genomic analyses comparing to adoptees. Non-adopted individuals were randomly placed into groups b, c and d.

### Phenotypic results

Phenotypic differences between adoptees and non-adopted individuals were generally modest in size but, due to the large sample size in this study, several were statistically significant (see Table 1). We found that non-adopted individuals showed significantly greater variance in their years of education than adoptees (26.2 vs 25.8; p=0.002 compared to 6500 non-adopted individuals in group d; p=3.2×10^−5^ compared to all non-adopted individuals). Table 1 gives descriptive statistics for education years, age and sex in the two groups. Adoptees in the UK Biobank were significantly younger on average (p=0.026 compared to group d; p=0.009 compared to all non-adopted individuals) although point estimates were similar (56.4 versus 56.7). There were significantly more males in the adopted group (p=0.033 compared to group d; p=0.008 compared to all non-adopted individuals), but the magnitude of the difference is small (48 versus 46% male). Adoptees had significantly fewer years of education (p=3.3×10^−11^ compared to group d; p< 2.2×10^−16^ compared to all non-adopted individuals in the UK Biobank). This is also reflected in the lower percentage of college attendees (20 years of education in Table 1) in the adopted group (28% compared to 33%). All comparison results were consistent between the large and small samples of non-adopted individuals.

**Table 1:** Comparative analysis of phenotypes in adoptees versus non-adopted individuals. Adoptees were compared to the full sample of non-adopted individuals, and to our smaller sub-sample used for genomic analyses (group d).

|  |  | Adopted (N=6311) | Non-adopted (N=375343) | Non-adopted (group d;<br>N=6500) |
| --- | --- | --- | --- | --- |
|  | Age | 56.4 (8.53) | 56.7 (8.01) | 56.7 (8.06) |
|  | Sex | 48% male (N=3010) | 46% male (N=172706) | 46% male (N=2978) |
| Education |  |  |  |  |
| Years | 7 | 1209 (19%) | 62651 (17%) | 1064 (16%) |
|  | 10 | 1780 (28%) | 100210 (27%) | 1709 (26%) |
|  | 13 | 749 (12%) | 43448 (12%) | 755 (12%) |
|  | 15 | 350 (6%) | 19428 (5%) | 354 (5%) |
|  | 19 | 433 (7%) | 24300 (6%) | 428 (7%) |
|  | 20 | 1790 (28%) | 125306 (33%) | 2190 (34%) |

### SNP heritability estimates

Figure 1 compares GREML-derived SNP heritability estimates for years of education in adopted individuals versus non-adopted individuals (left-hand bars). The estimate of heritability was larger in individuals reared with their relatives (0.29 [se = 0.079]) compared to adopted individuals (0.23 [se = 0.079]). However, confidence intervals were wide and overlapped, so the difference in heritability was not significant.

**Figure 1:**
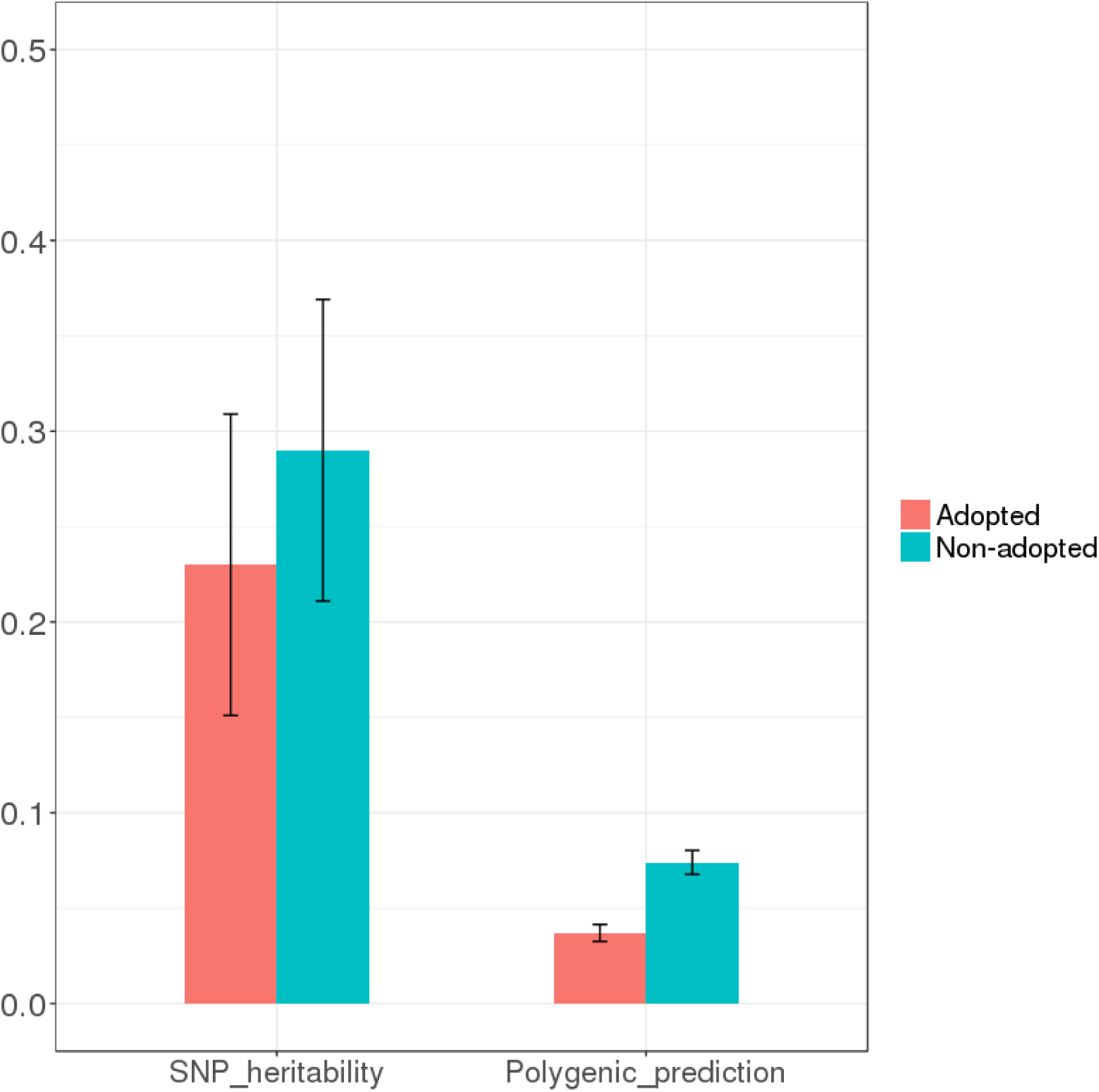
Estimates of the variance explained by common SNPs for years of education, and of the variance explained by polygenic scores for education polygenic scores, in adoptees compared to individuals who were reared with their relatives, plus 95% confidence intervals. Note: sample sizes for polygenic prediction analyses were 6,311 and 6,500 for adopted and non-adopted individuals respectively; sample sizes for GREML heritability analyses were lower (6,227 for adopted and 6,362 for non-adopted individuals) since relatives were removed at a cutoff of >0.025. For polygenic score results, CIs were obtained by bootstrapping with 1000 replications.

It was not computationally feasible to estimate the heritability of education using all non-adopted individuals with GREML. Notably, though, the LD score regression-derived heritability was 0.17 (se = 0.005) in the full sample of non-adopted individuals (N= 375,343), and 0.14 (se=0.073) for adoptees, corroborating the pattern of results found using GREML. LD score regression estimates are typically lower than GCTA-GREML-derived estimates (Evans and Keller 2018).

### Polygenic prediction results

Figure 1 also shows that twice as much phenotypic variance in years of education was explained by polygenic scores for education years in non-adopted individuals (0.074) as in adoptees (0.037). This difference was highly significant (p= 8.23 × 10^−24^). The optimal threshold for inclusion of SNPs was p=1 (Supplementary Table 1). Supplementary Table 2 shows the full results from the polygenic prediction analyses.

For our negative control analysis of height we found that, as expected, the variance explained by the polygenic score in adoptees (0.127, se = 0.008) versus non-adopted individuals (0.134, se = 0.008) was not significantly different (p=0.62). The optimal threshold for inclusion of SNPs in the polygenic score was p=0.001.

### Supplementary analyses

#### Heritability of being adopted

We found a liability scale SNP heritability of being adopted of 0.059 (se = 0.004), assuming the population prevalence of adoption is identical to the sample prevalence (1.7%). If the actual population prevalence differed and was, for example, 0.7% or 2.7%, the liability scale SNP heritability would become 0.047 (se = 0.002) or 0.066 (se = 0.005), respectively. Adoption status showed significant genetic correlations with education, age at first birth, depression and obesity after correcting for multiple testing (see Supplementary Table 5), although these correlations should be viewed with caution given the low SNP heritability of adoption. Adoption status could be significantly predicted by the education years polygenic score (R^2^=0.008, p<2 × 10^−16^). The heritability of adoption is low but may confound our between-group comparisons.

#### Polygenic score by adoption interaction

We tested a formal interaction model to further examine the finding that genetic influences on education are weaker in the sample of adoptees. The interaction between polygenic score and adoption status in predicting years of education is visualised in Figure 2. The regression slope is significantly steeper in the non-adopted group, indicating that years of education increases more as education polygenic scores increase in this group compared to the adopted group. See Supplementary Table 3 for the full interaction model results. Using linear regression, we confirmed that polygenic prediction of education interacts beyond additivity with adoption status (interaction estimate = −0.33; p=2.66 × 10^−4^). This means that polygenic scores had a smaller association with education in adoptees. Then using logistic regression instead of linear regression, we also found that the interaction exceeded multiplicativity (interaction estimate = −0.18; p = 0.0009). The finding of interaction exceeding both additive and multiplicative models means that the combined effect of education polygenic score and adoption status is not scale dependent and is greater than either the sum or product of their individual effects, respectively.

**Figure 2:**
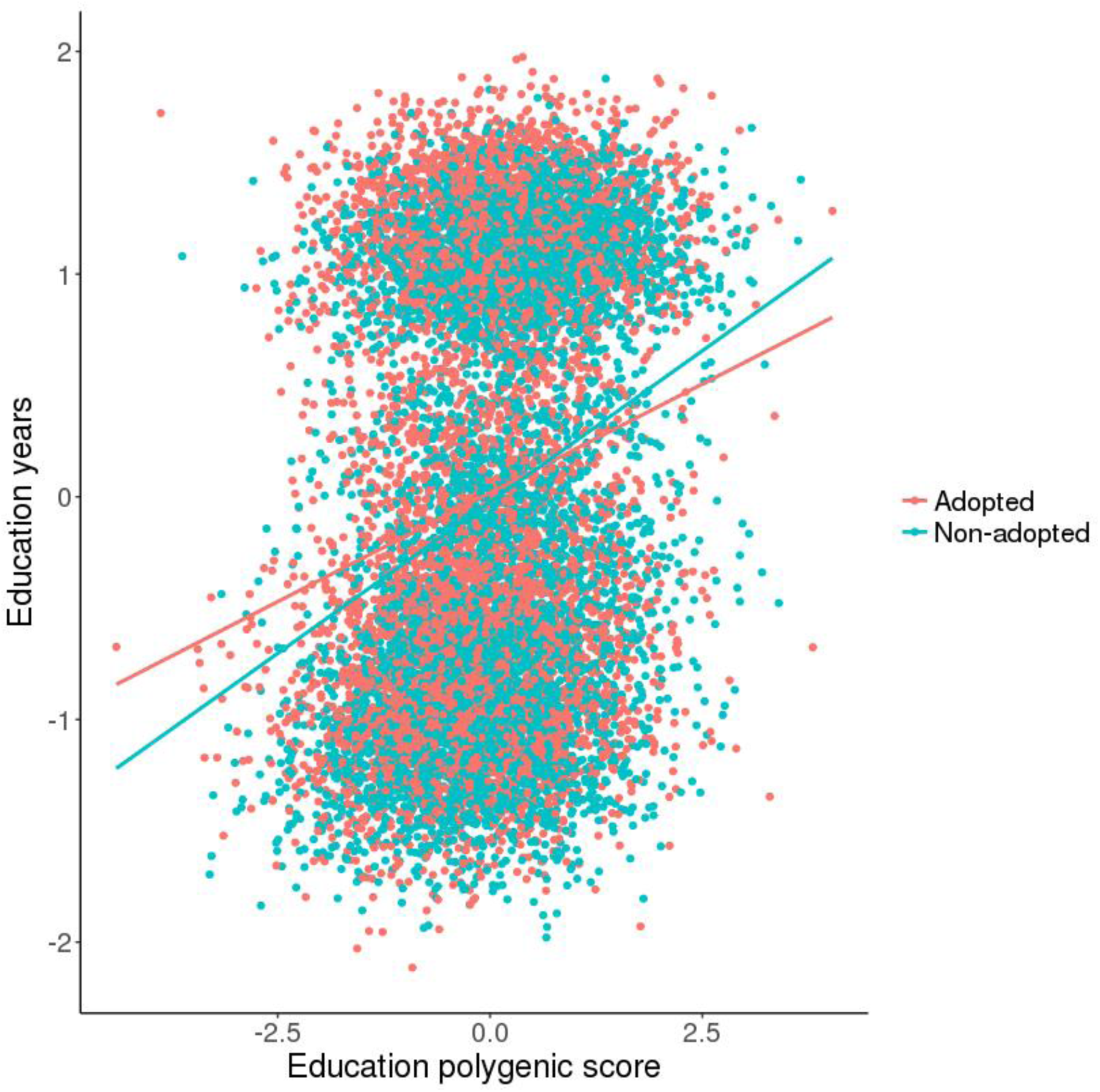
Regression of years of education on polygenic score for education, comparing 6311 adoptees to a sample of 6,500 non-adopted individuals. Note that the two clusters of data-points reflect the distinct groups of individuals who did and did not attend university.

To further explore the interaction, we plotted the mean years of education per decile of polygenic score for education years, for adopted and non-adopted individuals. Figure 3 shows that for individuals in the lowest decile of education polygenic score, those who were adopted as a child achieved a substantially higher mean years of education (standardised) compared to non-adopted individuals (−0.24 [se = 0.03] versus −0.40 [se = 0.03]). This mean difference between adopted and non-adopted individuals was significant in the bottom decile (p= 7.05×10^−5^), but not for other deciles of polygenic load. See Supplementary Table 4 for full results of the decile analysis.

**Figure 3:**
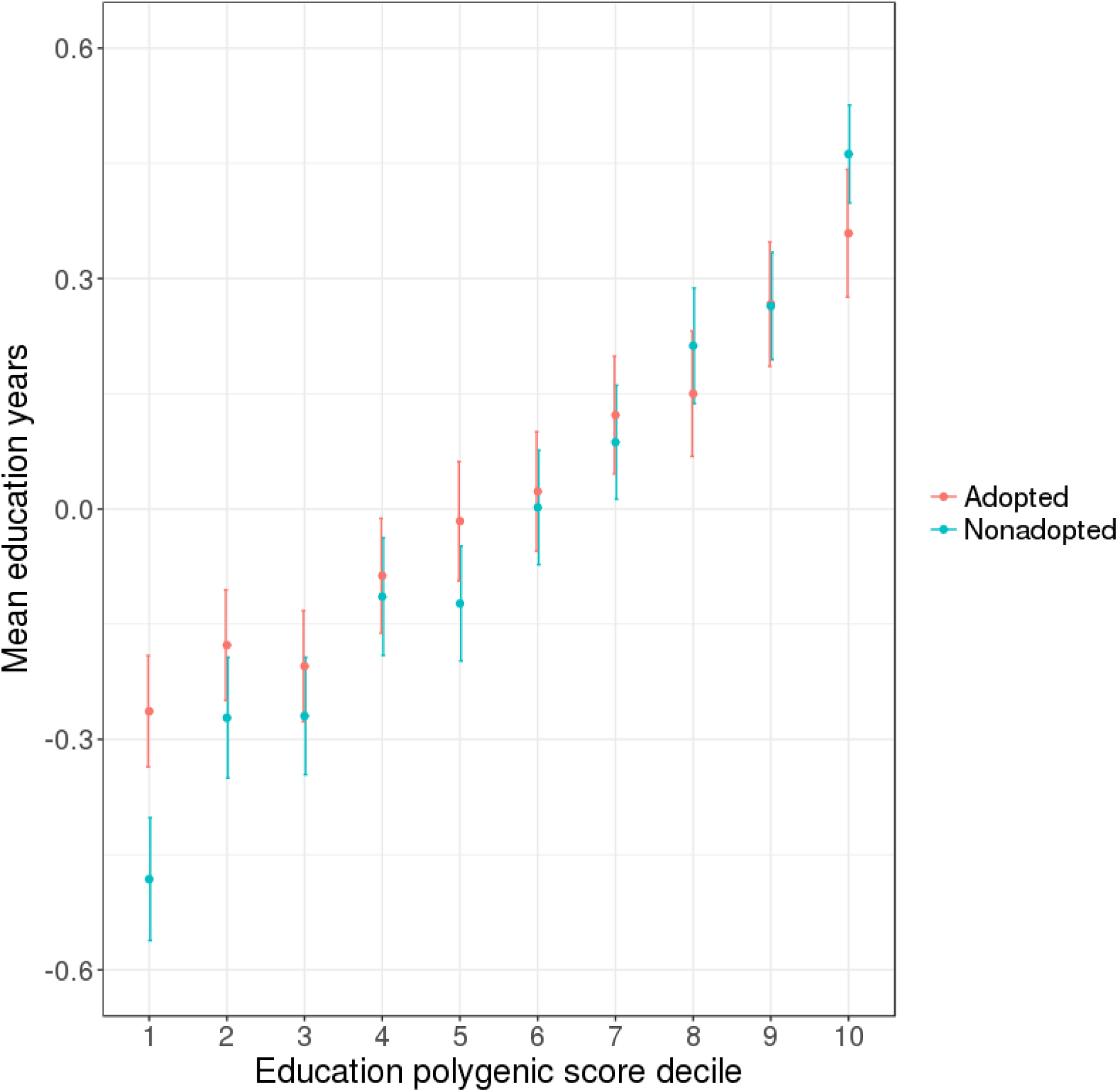
Mean education years (standardised) per decile of polygenic score for education years (also standardised), for adopted and non-adopted individuals, plus 95% confidence intervals. Mean education years differed significantly between adopted and non-adopted individuals in the lowest decile of education polygenic score, but not in others.

#### Qualitative differences in genetic influences according to adoptee status

We found that largely the same genetic influences are operating on education regardless of adoption status. First, the genetic correlation between education years in adopted and non-adopted individuals was not significantly different from 1 (0.81 [se = 0.21]). Second, we found no evidence that educational attainment is associated with different traits in individuals who were adopted. Supplementary Figure 1 presents estimates of genetic correlations between years of education and 247 external traits, comparing the adopted and non-adopted samples. None of these were significantly different between adoptees and non-adopted individuals after multiple testing correction. Due to the relatively small sample of adopted individuals, these results should be interpreted with caution.

#### Year-of-birth stratification analysis

Our final sensitivity analysis tested for differences according to year of birth in polygenic prediction from direct effects (indicated by the variance explained in the adoptees) versus from passive gene-environment correlation (indicated by the difference in variance explained between non-adopted individuals and adoptees). We found small, non-significant differences in the variance explained by polygenic scores for education depending on the year-of-birth group considered. Supplementary Figure 2 shows that polygenic prediction remains generally stable for the adoptees across generations at ∼0.04, and any differences between age strata were non-significant. We note that sub-sampling reduced the statistical power to detect differences within and between groups across time. See Supplementary Table 6 for sample sizes of each year-of-birth group.

## Discussion

Cumulatively, our findings suggest that the family environment provided by relatives plays an important role in the manifestation of genetic effects on education. The educational attainment of individuals who were adopted away from their parents as children had significantly less variance explained by polygenic scores (R^2^ =0.04 versus 0.07; difference test p= 8.23 × 10^−24^). The variance explained by polygenic scores in years of education in adoptees (0.04) can be regarded as an approximation of the prediction coming from the direct effects of individuals’ own DNA. The difference between the variance explained in non-adopted individuals and adoptees suggests that about half of the predictive power of polygenic scores for educational attainment comes from passive gene-environment correlation. In line with the difference in polygenic prediction between adoptees and non-adopted individuals, there was a significant and scale independent negative interaction, whereby adoption significantly decreases the polygenic prediction of education years. We further dissected this interaction by stratifying individuals by polygenic load for education, and found that adoptees in the lowest decile of polygenic score attaining significantly more years of education compared to those who were not adopted.

By showing that polygenic scores for education are twice as powerful in non-adopted individuals compared to adoptees, we suggest that genetic influence on educational attainment is magnified when individuals are reared by their close genetic relatives, with whom they share both genes and environments. Our results agree with recent evidence showing that the effects of passive gene-environment correlation reduced the variance explained by polygenic scores by 30-50% (Selzam et al. 2019; Kong et al. 2018). Notably, although we have controlled for the passive form by using adoptees, there are other gene-environment correlation mechanisms that are essential in how genes influence traits in everyone, including adoptees. These active and evocative processes (Plomin 2014) are part of what we term “direct genetic effects”.

Our observation that individuals in the lowest decile of education polygenic score attain significantly more education if they are adopted could be due to educationally supportive adoptive environments. This agrees with previous evidence showing that adopted individuals had higher school achievement and intelligence test scores than non-adopted siblings or peers who stayed with their birth family (van Ijzendoorn et al. 2005). Another adoption study previously found that adoptees performed better than non-adopted children from similar birth circumstances on childhood tests of reading, mathematics, and general ability, and retained this advantage in their later adult qualifications (Maughan et al. 1998). The environmental influence that adoptive parents appear to have on the attainment of individuals with low polygenic scores in this study might suggest that efforts to help individuals stay in education can be effective for those with less genetic propensity for education.

It is important to view these results in light of several limitations. First, interpreting genetic influence in adoptees as direct and free of passive gene-environment correlation requires that close biological relatives were not involved in the education of the adoptees. Unfortunately, the UK Biobank contains no information about the age of adoption beyond that it occurred in childhood, nor whether individuals were adopted by relatives. No information is collected on whether the adoption was ‘closed’ or ‘open’ – that is, whether individuals could identify and contact their biological parents. This knowledge would have allowed us to remove from our analyses individuals who were not solely socialised with adoptive families, and therefore to make a precise comparison to individuals who were reared with their birth parents. However, polygenic prediction of education still differed markedly between the two groups, even though adoptees may have been in contact with their biological relatives. Thus, the effects of passive gene-environment correlation may contribute even more than half of the predictive power of education polygenic scores, as we estimated here.

A second caveat is lack of generalisability. The UK Biobank is not representative of the general population, since there is apparent ‘healthy and wealthy’ volunteer selection (Keyes and Westreich 2019), and we have only analysed data on individuals with European ancestries. Furthermore, adoptive parents tend to differ systematically from other parents: they are likely to be more educated, more socially advantaged, and to have lower rates of psychopathology (Rutter 2006). This probably applies to our cohort, although we cannot be certain, due to the lack of parental data in the UK Biobank. If adoptive families are more homogeneous with respect to these characteristics, environmental variance may contribute less to differences in educational attainment among adoptees, and trait heritability estimates are consequently likely to be higher. However, the fact that lower environmental variance may act to *inflate* genetic influence in adoptees compared to non-adopted individuals makes our finding of significantly higher polygenic prediction in non-adopted individuals all the more striking. Again, the effects of passive gene-environment correlation for education may be even greater than we estimate.

There are several advantages of using the present adoption design for distinguishing direct genetic influence from passive gene-environment correlation. Unlike other methods, our approach does not require intergenerational data, which is valuable but has its own issues, such as cohort differences in genetic effects. Analysing the adoptees in the UK Biobank also bypasses several limitations of traditional adoption studies, including low sample size, and reliance on weak indirect proxies for inherited load for specific traits (birth parent trait status rather than individual-level polygenic scores). However, future progress in understanding the mechanisms driving the transmission of educational attainment will require intergenerational, longitudinal, genetically informative datasets, including detailed characterisation of the home environment. A developmental approach is helpful here, since gene-environment correlations likely arise early in childhood, when individuals interact closely with their relatives, and there will be complex reciprocal effects spanning through the life course. Researchers have already started to pinpoint genetically-influenced aspects of families that are associated with polygenic scores for education in the child generation (Wertz et al. 2018; Wertz et al. 2019; Krapohl et al. 2017).

The evidence presented in this study highlights the importance of the family environment to causal mechanisms influencing individual differences in educational attainment. These can be through possessing genes that shape the educational environment provided for offspring that also directly influence attainment in the child, or through providing an educationally supportive environment for your adopted child.

## Acknowledgments

We would like to thank the scientists involved in the construction of the UK Biobank and all of the participants who have shared their life experiences with investigators in the UK Biobank. This research has been conducted using the UK Biobank Resource, under the application 18177 (with thanks to Paul F. O’Reilly). This study represents independent research part funded by the National Institute for Health Research (NIHR) Biomedical Research Centre at South London and Maudsley NHS Foundation Trust and King’s College London. The views expressed are those of the author(s) and not necessarily those of the NHS, the NIHR or the Department of Health and Social Care. High performance computing facilities were funded with capital equipment grants from the GSTT Charity (TR130505) and Maudsley Charity (980). T.C. Eley is part funded by a program grant from the UK Medical Research Council (MR/M021475/1). RP is supported by a Medical Research Council Professorship award (G19/2). The positions of T.A. McAdams and Y.I. Ahmadzadeh, are funded by a Sir Henry Dale Fellowship awarded to T.A. McAdams, jointly funded by the Wellcome Trust and the Royal Society (grant number 107706/Z/15/Z). R. Cheesman is supported by an ESRC studentship. We thank Aysu Okbay and the SSGAC for providing genome-wide association summary statistics for educational attainment excluding UK Biobank.

## Additional information

Supplementary information is available for this paper.

