## Supplementary Information for "Comparison of adopted and non-adopted individuals reveals gene-environment interplay for education in the UK Biobank"

**Contents:**


- Supplementary Table 1: results from training education polygenic scores in 50,000 non-adopted individuals in the UK Biobank.
- Supplementary Table 2: full polygenic prediction results.
- Supplementary Table 3: full results from GxE (education polygenic score x adoption status) model.
- Supplementary Table 4: Comparison of adopted and non-adopted individuals for mean years of education per decile of polygenic score for education
- Supplementary Figure 1: genetic correlations between traits in LD Hub and years in education (plus standard errors) for adopted (red) and for non-adopted (blue) individuals.
- Supplementary Table 5: Genetic correlations with binary adopted/not adopted variable.
- Supplementary Figure 2: Comparing the variance explained in years of education by polygenic scores (PRS) for years of education, per year-of-birth grouping, for adoptees and non-adopted individuals.
- Supplementary Table 6: sample sizes by year-of-birth grouping.

*Supplementary Table 1: results from training education polygenic scores in 50,000 non-adopted individuals in the UK Biobank. Note: this p-value ($10^{-50}$) held for all of the standard set-values tested in PRSice (0.001, 0.05, 0.1, 0.2, 0.3, 0.4, 0.5, 1).*

| Threshold | PRS.$R^2$ | Full.$R^2$ | Null.$R^2$ | Coefficient | SE | P | Num_SNP |
|---|---|---|---|---|---|---|---|
| 1 | 0.071066 | 0.11043 | 0.039365 | 196340 | 3110.32 | $p < 10^{-50}$ | 87372 |

*Supplementary Table 2: full polygenic prediction results. PRS.SE = standard error around the $R^2$ value, derived from bootstrapping with 1000 replications. Note that the samples comprised 6500 non-adopted individuals and 6311 adoptees.*

| Group | PRS.$R^2$ | PRS.SE | Coefficient | SE | P | N-SNPs | Empirical P |
|---|---|---|---|---|---|---|---|
| Adopted | 0.037 | 0.0044 | 27581 | 1759.93 | $2.47\times10^{-54}$ | 87372 | $9.99\times10^{-05}$ |
| Non-adopted | 0.074 | 0.0063 | 38359.5 | 1681.36 | $6.78\times10^{-111}$ | 87372 | $9.99\times10^{-05}$ |

*Supplementary Table 3: full results from GxE (education polygenic score x adoption status) model. The outcome variable was years of education. All quantitative covariates were standardised. The table shows results for a linear model testing for additive interaction. The interaction term of interest is highlighted in yellow.*

| | Estimate | SE | t | Pr(>\|t\|) |
|---|---|---|---|---|
| (Intercept) | 13.42 | 0.4 | 33.91 | $<2x10^{-16}$ |
| PRS_std | 1.2 | 0.28 | 4.27 | $2x10^{-05}$ |
| adopted | 0.12 | 0.53 | 0.23 | 0.82 |
| Sex.0.0 | 0.63 | 0.12 | 5.24 | $1.66x10^{-07}$ |
| Age.at.recruitment.0.0_scaled | -0.8 | 0.06 | -12.8 | $<2x10^{-16}$ |
| PC1_scaled | 0.19 | 0.08 | 2.52 | 0.01 |
| PC2_scaled | -0.07 | 0.07 | -0.95 | 0.34 |
| PC3_scaled | 0.11 | 0.07 | 1.74 | 0.08 |
| PC4_scaled | $4.1x10^{-03}$ | 0.06 | 0.07 | 0.95 |
| PC5_scaled | -0.03 | 0.07 | -0.5 | 0.62 |
| PC6_scaled | -0.13 | 0.07 | -1.92 | 0.06 |
| PC7_scaled | -0.07 | 0.07 | -1.1 | 0.27 |
| PC8_scaled | -0.05 | 0.06 | -0.87 | 0.38 |
| PC9_scaled | 0.12 | 0.08 | 1.53 | 0.13 |
| PC10_scaled | -0.22 | 0.07 | -3.15 | $1.61x10^{-03}$ |
| Centre11002 | 0.69 | 0.52 | 1.31 | 0.19 |
| Centre11003 | 0.11 | 0.52 | 0.21 | 0.83 |
| Centre11004 | -0.27 | 0.51 | -0.53 | 0.59 |
| Centre11005 | 1.15 | 0.52 | 2.2 | 0.03 |
| Centre11006 | 0.03 | 0.5 | 0.07 | 0.95 |
| Centre11007 | 1.21 | 0.47 | 2.6 | 0.01 |
| Centre11008 | -0.14 | 0.46 | -0.3 | 0.76 |
| Centre11009 | -0.56 | 0.47 | -1.2 | 0.23 |
| Centre11010 | 0.17 | 0.44 | 0.39 | 0.7 |
| Centre11011 | 0.2 | 0.45 | 0.44 | 0.66 |

| | | | | |
|---|---|---|---|---|
| Centre11012 | 1.54 | 0.57 | 2.69 | 0.01 |
| Centre11013 | 0.36 | 0.46 | 0.78 | 0.43 |
| Centre11014 | -0.22 | 0.46 | -0.49 | 0.63 |
| Centre11016 | -0.02 | 0.45 | -0.05 | 0.96 |
| Centre11017 | -0.09 | 0.48 | -0.18 | 0.86 |
| Centre11018 | 1.07 | 0.48 | 2.22 | 0.03 |
| Centre11020 | 0.64 | 0.48 | 1.33 | 0.18 |
| Centre11021 | 0.48 | 0.47 | 1.01 | 0.31 |
| Centre11022 | -0.62 | 0.95 | -0.65 | 0.52 |
| Centre11023 | -3.52 | 1.51 | -2.33 | 0.02 |
| PRS_std:adopted | -0.33 | 0.09 | -3.65 | $2.66 \times 10^{-04}$ |
| PRS_std:Sex.0.0 | -0.06 | 0.09 | -0.66 | 0.51 |
| PRS_std:Age.at.recruitment.0.0_scaled | -0.03 | 0.04 | -0.64 | 0.52 |
| PRS_std:PC1_scaled | -0.04 | 0.05 | -0.93 | 0.35 |
| PRS_std:PC2_scaled | -0.03 | 0.05 | -0.69 | 0.49 |
| PRS_std:PC3_scaled | -0.1 | 0.04 | -2.21 | 0.03 |
| PRS_std:PC4_scaled | -0.05 | 0.04 | -1.08 | 0.28 |
| PRS_std:PC5_scaled | -0.01 | 0.05 | -0.25 | 0.80 |
| PRS_std:PC6_scaled | 0.03 | 0.05 | 0.72 | 0.47 |
| PRS_std:PC7_scaled | 0.02 | 0.05 | 0.38 | 0.70 |
| PRS_std:PC8_scaled | 0.02 | 0.04 | 0.37 | 0.71 |
| PRS_std:PC9_scaled | $1.7 \times 10^{-04}$ | 0.05 | $4 \times 10^{-03}$ | 1.00 |
| PRS_std:PC10_scaled | $3.6 \times 10^{-03}$ | 0.05 | 0.08 | 0.94 |
| PRS_std:Centre11002 | 0.37 | 0.36 | 1.03 | 0.30 |
| PRS_std:Centre11003 | 0.46 | 0.36 | 1.28 | 0.20 |
| PRS_std:Centre11004 | -0.27 | 0.37 | -0.74 | 0.46 |
| PRS_std:Centre11005 | 0.76 | 0.37 | 2.09 | 0.04 |
| PRS_std:Centre11006 | 0.25 | 0.35 | 0.73 | 0.47 |

| | | | | |
|---|---|---|---|---|
| PRS_std:Centre11007 | 0.25 | 0.33 | 0.75 | 0.45 |
| PRS_std:Centre11008 | 0.01 | 0.33 | 0.04 | 0.97 |
| PRS_std:Centre11009 | 0.14 | 0.33 | 0.43 | 0.67 |
| PRS_std:Centre11010 | -0.14 | 0.31 | -0.45 | 0.65 |
| PRS_std:Centre11011 | 0.08 | 0.31 | 0.26 | 0.8 |
| PRS_std:Centre11012 | 0.45 | 0.39 | 1.16 | 0.25 |
| PRS_std:Centre11013 | 0.03 | 0.32 | 0.11 | 0.92 |
| PRS_std:Centre11014 | 0.19 | 0.33 | 0.59 | 0.56 |
| PRS_std:Centre11016 | 0.26 | 0.32 | 0.8 | 0.43 |
| PRS_std:Centre11017 | 0.11 | 0.35 | 0.33 | 0.74 |
| PRS_std:Centre11018 | 0.35 | 0.34 | 1.03 | 0.31 |
| PRS_std:Centre11020 | 0.39 | 0.34 | 1.16 | 0.24 |
| PRS_std:Centre11021 | 0.21 | 0.33 | 0.63 | 0.53 |
| PRS_std:Centre11022 | -0.21 | 0.62 | -0.34 | 0.73 |
| PRS_std:Centre11023 | 0.14 | 1.03 | 0.14 | 0.89 |
| adopted:Sex.0.0 | -0.27 | 0.17 | -1.58 | 0.12 |
| adopted:Age.at.recruitment.0.0_scaled | -0.23 | 0.09 | -2.70 | 0.01 |
| adopted:PC1_scaled | -0.03 | 0.09 | -0.37 | 0.71 |
| adopted:PC2_scaled | 0.04 | 0.10 | 0.46 | 0.64 |
| adopted:PC3_scaled | 0.03 | 0.09 | 0.36 | 0.72 |
| adopted:PC4_scaled | -0.07 | 0.09 | -0.85 | 0.39 |
| adopted:PC5_scaled | 0.11 | 0.09 | 1.16 | 0.25 |
| adopted:PC6_scaled | 0.05 | 0.10 | 0.49 | 0.62 |
| adopted:PC7_scaled | -0.02 | 0.10 | -0.25 | 0.80 |
| adopted:PC8_scaled | 0.12 | 0.09 | 1.40 | 0.16 |
| adopted:PC9_scaled | -0.15 | 0.09 | -1.60 | 0.11 |
| adopted:PC10_scaled | 0.12 | 0.09 | 1.28 | 0.20 |
| adopted:Centre11002 | -0.44 | 0.72 | -0.61 | 0.54 |

| | | | | |
|---|---|---|---|---|
| adopted:Centre11003 | -0.61 | 0.70 | -0.87 | 0.39 |
| adopted:Centre11004 | 0.10 | 0.71 | 0.13 | 0.89 |
| adopted:Centre11005 | -1.12 | 0.73 | -1.52 | 0.13 |
| adopted:Centre11006 | 0.07 | 0.68 | 0.10 | 0.92 |
| adopted:Centre11007 | -1.42 | 0.64 | -2.23 | 0.03 |
| adopted:Centre11008 | -0.03 | 0.63 | -0.06 | 0.96 |
| adopted:Centre11009 | 0.27 | 0.63 | 0.43 | 0.67 |
| adopted:Centre11010 | -0.68 | 0.59 | -1.14 | 0.26 |
| adopted:Centre11011 | -0.42 | 0.60 | -0.70 | 0.48 |
| adopted:Centre11012 | -0.73 | 0.77 | -0.95 | 0.34 |
| adopted:Centre11013 | -0.88 | 0.62 | -1.43 | 0.15 |
| adopted:Centre11014 | 0.07 | 0.64 | 0.12 | 0.91 |
| adopted:Centre11016 | -0.29 | 0.62 | -0.46 | 0.64 |
| adopted:Centre11017 | 0.02 | 0.67 | 0.03 | 0.97 |
| adopted:Centre11018 | -0.23 | 0.66 | -0.35 | 0.73 |
| adopted:Centre11020 | -0.59 | 0.66 | -0.91 | 0.36 |
| adopted:Centre11021 | -0.53 | 0.65 | -0.82 | 0.41 |
| adopted:Centre11022 | 0.96 | 1.29 | 0.74 | 0.46 |
| adopted:Centre11023 | 1.04 | 2.15 | 0.48 | 0.63 |

*Supplementary Table 4: Comparison of adopted and non-adopted individuals for mean years of education per decile of polygenic score for education.*

| | Adopted | | | non-adopted | | | | |
|---|---|---|---|---|---|---|---|---|
| Decile | Mean education years | SE | N | Mean education years | SE | N | zdiff | zdiffp |
| 1 | -0.26 | 0.04 | 723 | -0.48 | 0.04 | 559 | -3.97 | **$7.07 \times 10^{-05}$** |
| 2 | -0.18 | 0.04 | 710 | -0.27 | 0.04 | 571 | -1.74 | 0.081 |
| 3 | -0.20 | 0.04 | 672 | -0.27 | 0.04 | 609 | -1.21 | 0.225 |
| 4 | -0.09 | 0.04 | 640 | -0.11 | 0.04 | 641 | -0.50 | 0.619 |
| 5 | -0.02 | 0.04 | 638 | -0.12 | 0.04 | 643 | -1.95 | 0.051 |
| 6 | 0.02 | 0.04 | 623 | 0.002 | 0.04 | 658 | -0.37 | 0.708 |
| 7 | 0.12 | 0.04 | 625 | 0.09 | 0.04 | 656 | -0.65 | 0.515 |
| 8 | 0.15 | 0.04 | 583 | 0.21 | 0.04 | 698 | 1.11 | 0.269 |
| 9 | 0.27 | 0.04 | 562 | 0.26 | 0.04 | 719 | -0.05 | 0.962 |
| 10 | 0.36 | 0.04 | 535 | 0.46 | 0.03 | 746 | 1.93 | 0.053 |

*Supplementary Figure 1: genetic correlations between traits in the LD Hub database and years of education (plus standard errors) for adopted (red) and for non-adopted (blue) individuals. The figure only shows results for traits that showed significant genetic correlations with education in the sample of adoptees at p<0.05.*

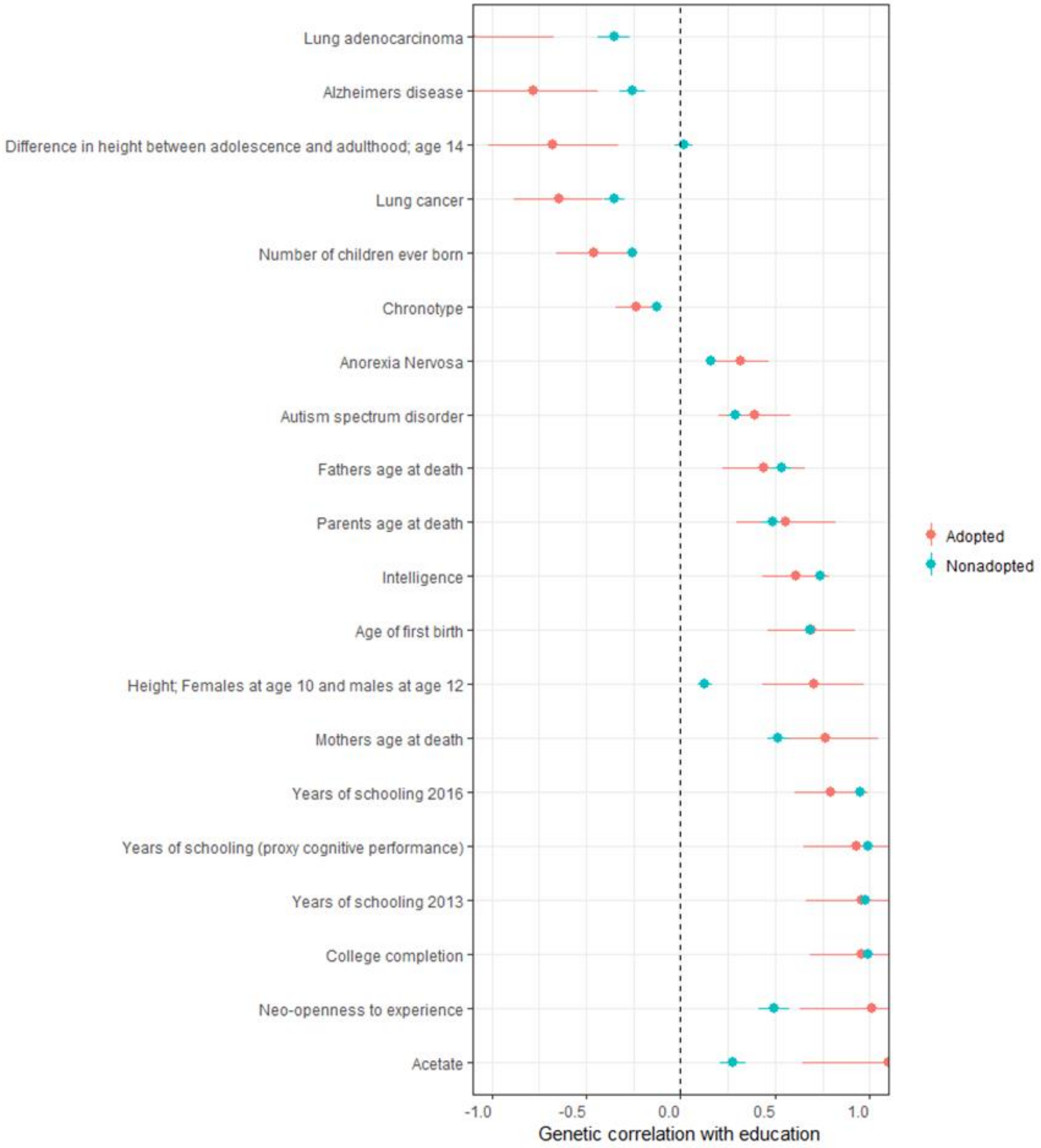


Due to the relatively small sample of adoptees, only 4 traits were significantly genetically correlated with education years in the adopted sample after correcting for multiple testing by using a stringent p-value of 0.000202 (0.05/247): years of schooling (2016), college completion, intelligence, and years of schooling (proxy). All traits in this figure also showed significant genetic correlations with education years in the non-adopted sample at p<0.000202, except height of females age 10 and males at age 12 (p=0.0004), Alzheimer's disease (p=0.0002) and difference in height between adolescence and adulthood (p=0.7265). None of the genetic correlations with education were significantly different between adoptees and non-adopted individuals at p<0.000202.

*Supplementary Table 5: Genetic correlations with binary adopted/not adopted variable, from LD Hub, ranked by p-value. The table only shows correlations that remained significant after a Bonferroni correction for multiple testing (p< 0.0002 i.e. 0.05/247).*

| Trait | Genetic correlation | se | p |
|---|---|---|---|
| Years of schooling 2016 | -0.5216 | 0.0615 | $2.28x10^{-17}$ |
| Age of first birth | -0.6589 | 0.0836 | $3.23x10^{-15}$ |
| College completion | -0.5584 | 0.0882 | $2.41x10^{-10}$ |
| Years of schooling (proxy cognitive performance) | -0.522 | 0.0851 | $8.38x10^{-10}$ |
| Years of schooling 2013 | -0.4739 | 0.0823 | $8.39x10^{-09}$ |
| Depressive symptoms | 0.5176 | 0.1026 | $4.60x10^{-07}$ |
| Intelligence | -0.306 | 0.0679 | $6.66x10^{-06}$ |
| Obesity class 1 | 0.2796 | 0.0652 | $1.77x10^{-05}$ |
| Waist-to-hip ratio | 0.2507 | 0.0626 | $6.23x10^{-05}$ |
| Obesity class 2 | 0.3368 | 0.0851 | $7.58x10^{-05}$ |
| Extreme bmi | 0.2943 | 0.0748 | $8.32x10^{-05}$ |
| Mothers age at death | -0.6567 | 0.1679 | $9.14x10^{-05}$ |
| Body mass index | 0.2486 | 0.0648 | 0.0001 |
| Insomnia | 0.4258 | 0.1098 | 0.0001 |

Note: These significant negative genetic correlations with educational attainment, age of motherhood, and poor physical health agree with evidence that adoptees are more likely than individuals from comparable origins to be born to young mothers, to have had a low birthweight, and to have received suboptimal obstetric care (Maughan et al. 1998), although the two groups were highly similar apart from these factors. The fact that adoptee status had a low heritability (6%; liability scale) means that interpretation of these genetic correlations should be cautious.

*Supplementary Figure 2: Comparing the variance explained in years of education by polygenic scores (PRS) for years of education, per year-of-birth grouping, for adoptees and non-adopted individuals. 95% CIs were obtained by bootstrapping with 1000 replications.*

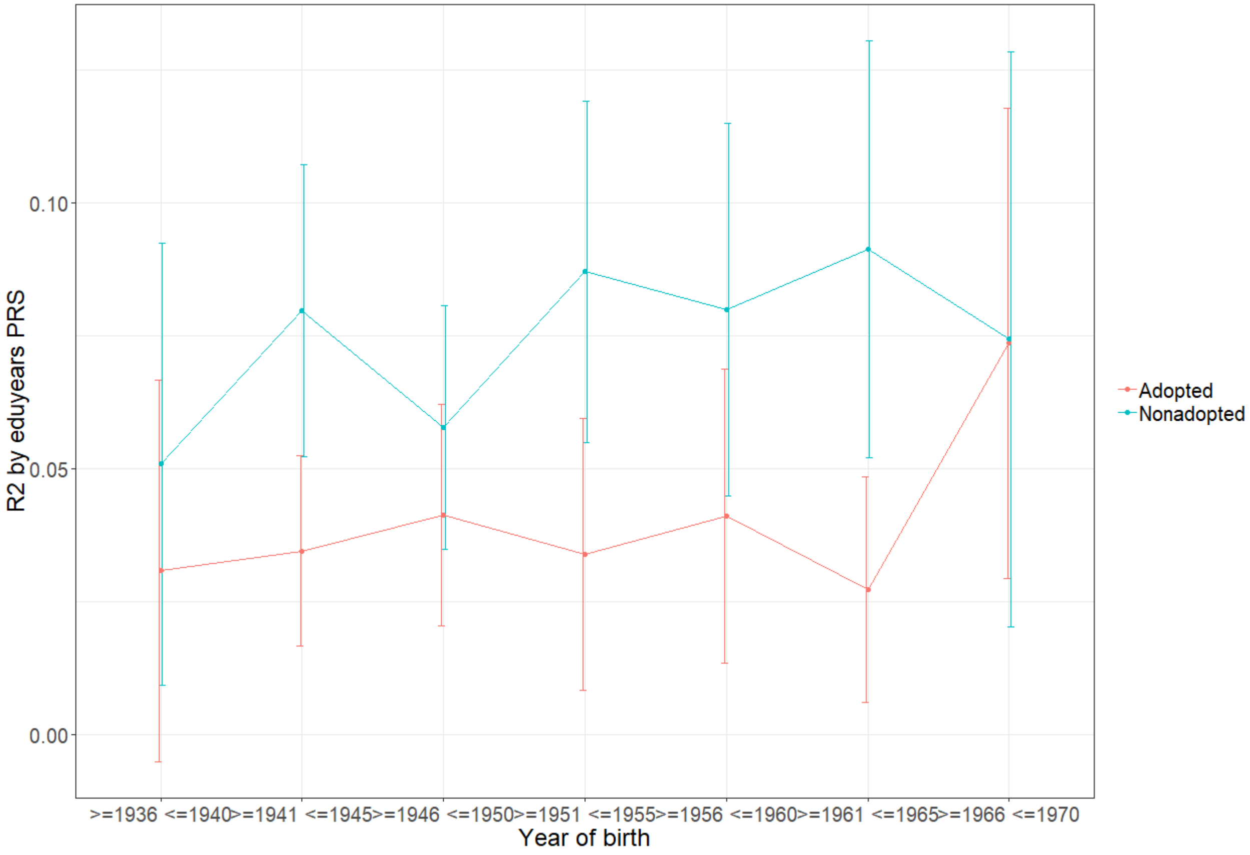


Any systematic differences between adoptive and biological families are likely to have changed over time. The variance explained by polygenic scores is more variable in the group of individuals reared by their biological relatives, but generally increases over time. The plot shows that prediction is stronger for adoptees in the youngest age group (although this increase was not significant). Due to increasing use of contraceptives across the 20th century, adoption became less related to young motherhood and more related to removal of children from high risk environments. Adoptions occurring later in the twentieth century were more likely to involve older children rather than babies (Maughan et al. 1998; Keating 2008). We speculate that these changes could mean that younger adoptees in our sample are less generalisable, more likely to have come from high risk environments, and more likely to have spent more time with their biological relatives, experiencing passive gene-environment correlation. However, other than the non-significant spike in the youngest age group of adoptees, polygenic prediction remains stable across cohorts (at ~4%). It could be argued that the variance explained by polygenic scores in adoptees provide a rough benchmark of 'direct' genetic influence remaining throughout periods of great social change.

*Supplementary Table 6: sample sizes by year-of-birth grouping.*

| | Adopted | Non-adopted group d (N=6500) |
|---|---|---|
| >=1936 <=1940 | 341 | 448 |
| >=1941 <=1945 | 1687 | 1413 |
| >=1946 <=1950 | 1312 | 1503 |
| >=1951 <=1955 | 839 | 1124 |
| >=1956 <=1960 | 781 | 867 |
| >=1961 <=1965 | 869 | 796 |
| >=1966 <=1970 | 482 | 349 |

The post-war spike in the number of adoptions (and number of births generally) is consistent with the ‘baby scoop’ and the ‘baby boom’.
